# The ModelSEED Biochemistry Database, 2026 update: grading multi-source thermodynamics

**DOI:** 10.64898/2026.09.15.751859

**Authors:** Andrew Freiburger, José P. Faria, Ray Zhu, Janaka N. Edirisinghe, Filipe Liu, Cooper Taylor, Vibhav Setlur, Robert T. Giessmann, Moritz E. Beber, Elad Noor, Vikas Upadhyay, Mohit Anand, Costas D. Maranas, Sebastian Huß, Zoran Nikoloski, Adam P. Arkin, Robert W. Cottingham, Elisha M. Wood-Charlson, Arvind Ramanathan, Christopher S. Henry, Samuel M. D. Seaver

## Abstract

The ModelSEED Biochemistry Database (https://modelseed.org/biochem) supplies foundational mass- and charge-balanced reaction networks for metabolic reconstructions. Here we present an update on the biochemistry where the database has been expanded to ∼ 46, 000 compounds, featuring ∼37, 000 metabolic structures, and ∼48, 000 reactions. We expanded our approach for handling thermodynamic data, enabling multiple sources of data to be derived, integrated, and presented to the wider research community. We now publish predictions of pKa, reaction energy (and respective uncertainties), and estimates of reaction direction from multiple independent sources. Reaction directions are graded gold, silver, or bronze according to the strength of supporting evidence, which is complemented by the prediction from an ensemble of large language models. These diverse sources empower users to understand agreements or discrepancies in directionality for ∼27, 000 reactions. Finally, to ensure data integrity, a new conflict-resolution pipeline reconciles structures across sources, documenting input from curators. Our work is publicly available at https://github.com/ModelSEED/ModelSEEDDatabase.

## Introduction

The ModelSEED Biochemistry Database provides mass- and charge-balanced reaction networks reconciled across major biochemical resources for metabolic reconstructions (4). Since its 2020 release, the database has been widely adopted for curating genome-scale metabolic models (GEMs): e.g. GEMsembler utilizes it for cross-mapping between resources (5) and Yeast9 sources Gibbs energies from it for constraining the core GEM (6). This update reports an expansion of the database, a tiered integration of thermodynamic sources spanning heuristic methods to large language models, and graded confidence scores for reaction directionality based on supporting evidence. While > 70% of sources agree on reaction directionality, disagreements are displayed for investigators to interpret. Atom-mapping data is further integrated to enable metabolic flux analysis (MFA), and a structure-curation pipeline is built to resolve conflicts between structures from different databases. Finally, we expanded our search interface at https://modelseed.org/biochem to facilitate user queries of the resource.

## Materials and Methods

### Construction and Curation

We collated additional compounds and reactions from MetaCyc (7), Rhea (3), and ChEBI (2), merging compounds via structures where possible and matching reactions by compound identities, similar to the 2020 release (4). Mass and charge balances are enforced and flagged. Unlike the 2020 release, where compounds with conflicting sourced structures were either manually reviewed or silently dropped, the updated pipeline classifies structural conflicts and either 1) automatically picks from the source that matches the historical formula, 2) prioritizes curator authored and justified content, or 3) excludes (with logged justification) compounds without a defensible source formula. This enforces that every outcome is attributed to its curator and evidence. Every structure is retained in the repository to facilitate conflict resolution. Investigators can contribute to the ModelSEED Database through the procedure in Supplementary Methods S1, which is expanded in the repository.

### Multi-source thermodynamics

The 2020 release combined two thermodynamics sources: eQuilibrator (8) and the group contribution method (9). This ModelSEED Database update incorporates the most recent eQuilibrator 3.0 (10) and introduces two new thermodynamics sources: dGPredictor (11) and experimental values. Each of these four distinct thermodynamics sources are reported separately where available. All values are reported at pH 7.0, ionic strength 0.25 M, pMg 3.0 and 298.15 K; the basis for each condition and its effect on the released energies are given in Supplementary Methods S2.

#### Protonation

eQuilibrator’s Legendre transform requires a *macroscopic* pKa ladder per compound: an ordered list in which consecutive species differ by one proton. The count of entries around the desired pH sets the proton count of the major microspecies, so the composition of the ladder changes the transformed energy directly. We generate the pKa ladders for every compound with a structure (source prevenance preserved), and the measured ladders of Alberty (12), using licensed ChemAxon Marvin 26.1 (13) for use with eQuilibrator and also via open-source MolGpKa (14) for unlicensed use.

#### Experimental anchors

eQuilibrator and dGPredictor draw on the same set of experimental values derived from the NIST Thermodynamics of Enzyme-catalyzed Reactions (TECR) Database (15). There were structural conflicts in compounds mapped to ModelSEED from the KEGG (1) IDs of TECR and the MetaNetX (16) IDs of eQuilibrator, so to facilitate direct comparisons between thermodynamic predictions of ModelSEED reactions, we re-built the experimental anchors of TECR values and curated conflicts to map TECR directly to ModelSEED. We further investigated openTECR, a communal re-curation of TECR available at https://github.com/opentecr, to extend the thermodynamic anchors.

#### Direction

To conserve precedence, we use the same 2020 heuristics to derive reaction directions from the Group contribution approach (9), so that series stays comparable across releases. For eQuilibrator and dGPredictor, we use one rule set built on the reversibility index from Noor *et al*. (17): irreversible when | ln Γ| exceeds ln(1000) by more than one propagated standard deviation (10), where Γ is the fold-change of reactant concentration required to change reaction direction. We extend the rule to require that the uncertainty clears this Γ threshold as well; however, differences in how each source estimates its uncertainty prevent direct comparisons (Figure 2C)).

### Grading reaction direction

We apply a classification approach with which we grade the predicted reaction direction based on the quality of available evidence. The selected tiers are intuitively described as gold (highest), silver (intermediate), or bronze (lowest). Our evaluation splits two ways: 1) confidence of a source’s own claim fit as a probability against the experimental anchors and 2) a weighted comparison of other sources on the same scale. Grades, self-assessments, and cross-source verdicts are stored in the biochemistry database and exposed on the website for users inspection. The grading methodology is thoroughly described in Supplementary Methods S3.

### Ensemble LLMs predictions

Thermodynamic assignment is bounded by estimate coverage, which means that energies and therefore directions cannot be computed for 23,790 structurally incomplete reactions. We therefore run a complementary approach using an ensemble or “council” of large language models (LLMs) to estimate directionality from only reaction name and stoichiometry, where three LLM agents independently predict a direction, a fourth audits, and a fifth adjudicates. We provide the prompts in the repository, and describe the process and its limitations in Supplementary Methods S4.

### Atom mapping

Atom mappings are generated from the workflow of Huß *et al*. (18) where they refine the per-reaction output of the Reaction Decoder Tool (19) into mappings that are comprehensive enough for reconstructing a metabolic model. The approach also resolves chemically equivalent atoms into symmetry groups, such as the two oxygens of CO_2_ and the six equivalent terminal oxygens of pyrophosphate, so the ambiguity is apparent. The output is formatted and stored in the database, and can also be visualized in the UI.

### Distribution and community process

The database is distributed as versioned flat files and JSON from the public repository, queried programmatically through a SOLR endpoint, and browsed through the web interface at https://modelseed.org/biochem. Every merge runs continuous integration that rebuilds the database from source records and fails on any regression in balance, structure coverage, or schema conformance to prevent curation changes from breaking the database. Anyone can contribute by adding and curating structures and reactions, following guidance in the repository via pull requests.

## Results

### Growth and coverage

The database has grown 34% in non-obsolete compounds and reactions since 2020, driven mainly by MetaCyc (7) and the BioCyc family (20). Reaction growth is now split across three primary databases (Figure 1B): MetaCyc (7) carries 26,140 reactions of which 18,893 are unique among the primary sources, KEGG (1) 10,840 of which 5,291 are unique, and the newly-integrated Rhea (3) 14,511 of which 8,210 are unique. The completeness of reactions, meaning whether all involved compounds are structurally defined, differs among (Figure 1C): 90% for MetaCyc, 75% for KEGG, 65% for Rhea. Overall structure coverage rose from 28,087 to 36,907 compounds, with the distribution of this new biochemistry described in Supplementary Methods S5 and Supplementary Figure S1.

**Figure 1.**
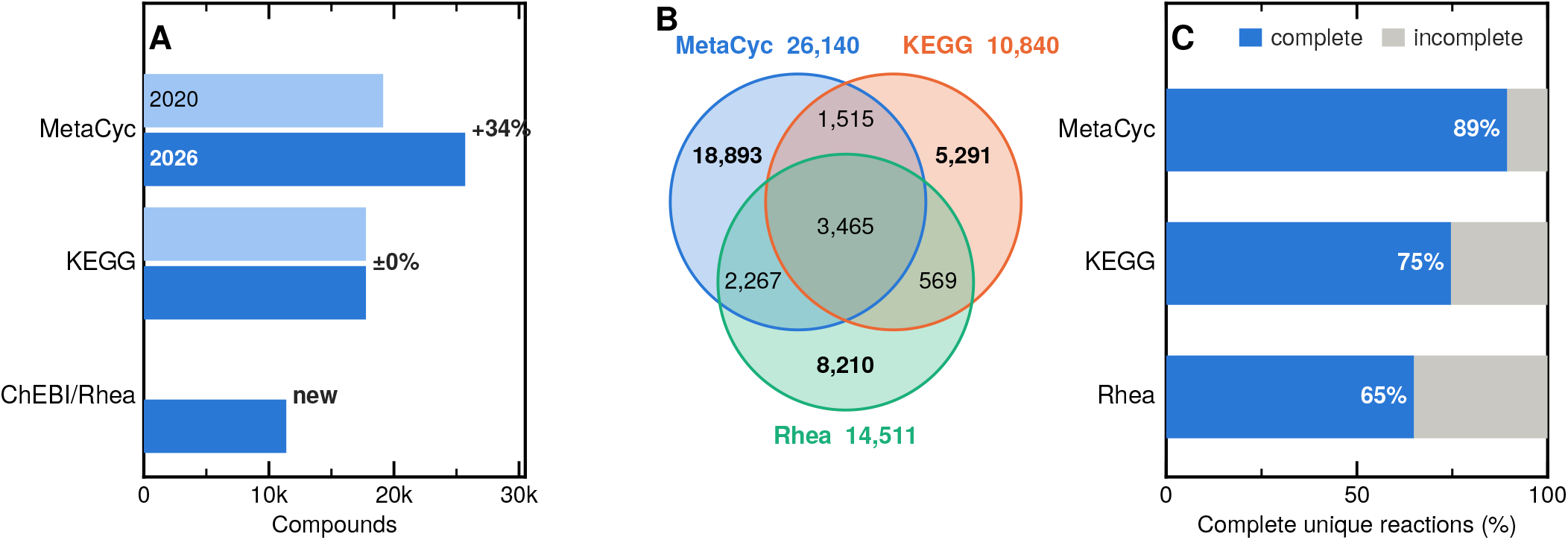
Sources of compounds and reactions. (**A**) compounds contributed by each source that also supplied molecular structures, 2020 against 2026: we did not introduce any new biochemistry from KEGG (1). The third row is ChEBI (2), which supplies structures but no reactions. (**B**) how the three primary reaction sources overlap, every region labelled with its reaction count. Circle areas are not proportional. Rhea (3) identifies its compounds through ChEBI. (**C**) the share of each source’s unique reactions that are complete, meaning every reagent is assigned a complete structure, so the reaction can be balanced and decomposed.

**Figure 2.**
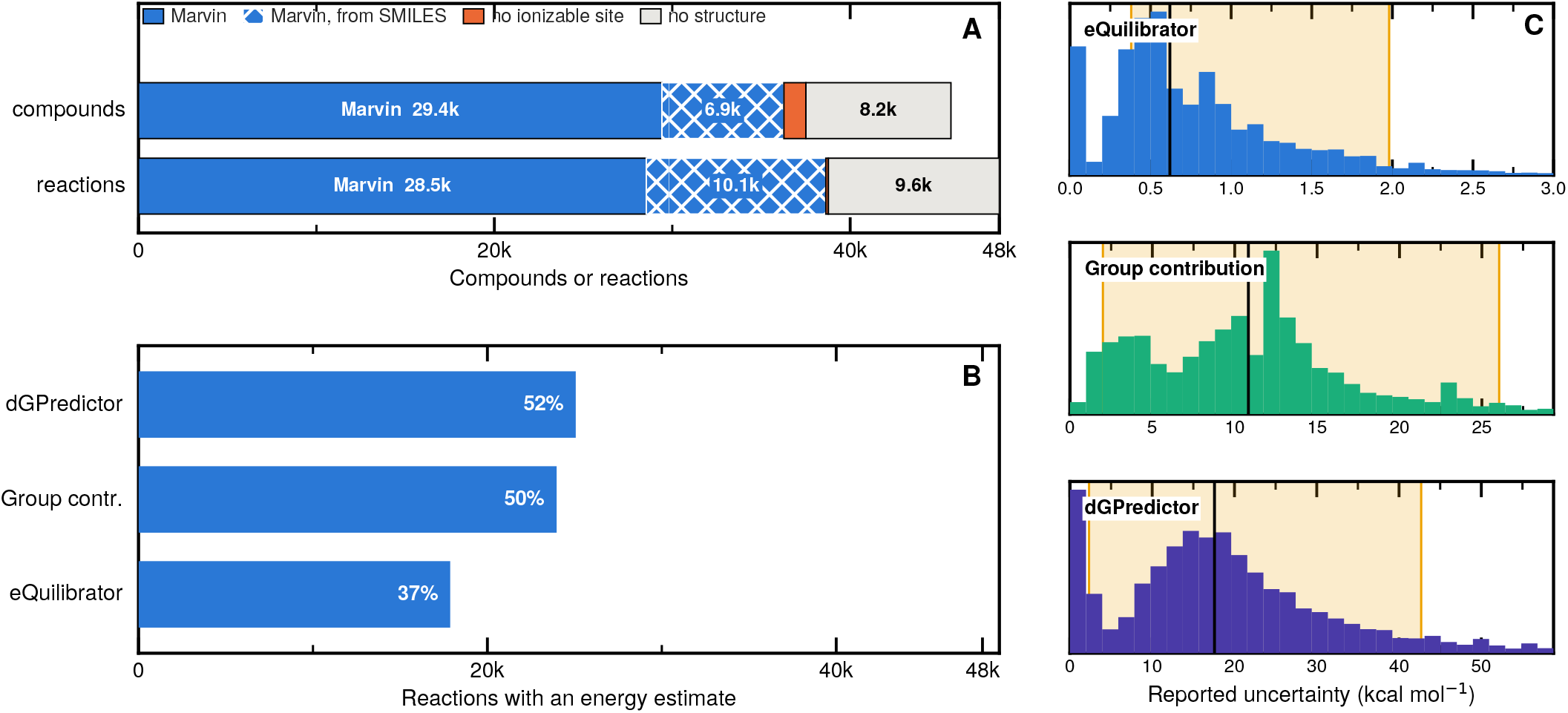
Thermodynamic sources. (**A**) Provenance of every pK_a_ ladder shipped in the database, across all 45,662 compounds and 48,384 reactions. ChemAxon Marvin supplies 79.4% of compounds and 79.8% of reactions. *Hatching* marks the share computed from SMILES because Marvin refuses polymers and organometallics as query molecules. *No ionizable site* edscribes 1,237 compounds that lacked a dissociable proton from Marvin between pH −2 and 16. (**B**) Reactions that carry an energy computed from the source heuristic. (**C**) Reported uncertainties (kcal mol^−1^) for each source, where the shaded band spans the 5th to 95th percentile range used in our classification scheme.

### Structure curation

We built and ran our curation pipeline on all structures that would impact a prioritized set of roughly 9,000 reactions used in ModelSEED (21; 22) and PlantSEED (23; 24). This scope is not a statement about how much of the database is usable. The conflict pipeline resolved every previously conflicting compound in a three-staged curation: i) historical formula were used to automatically resolve conflicts in mobile hydrogens, affecting 60 compounds and 550 reactions; ii) manual curation resolved conflicts in molecular structures, affecting 46 compounds and 59 reactions; and iii) where steps i-ii fail because of an absence in defensible source formula, reactions were added to a mass-balance exclusion registry where we fix an arbitrarily chosen formula, affecting 4 compounds and 61 reactions.

### Thermodynamics

The three predictors cover comparable fractions of the database and disagree substantially about how well they do it (Figure 2B).

Uncertainty is reported on scales differing by more than an order of magnitude (Figure 2C): median errors (kcal mol^−1^) of 0.62 for eQuilibrator, 10.83 for group contribution, and 17.57 for dGPredictor.

#### Calibrating the reported errors against measurement

We use the experimental anchors to evaluate the error estimates. For each source, we collect one held-out prediction per anchored reaction, together with the uncertainty that source reports for it, and divide the residual against the measured Δ_r_*G*^*′°*^ by that uncertainty. Measured this way, eQuilibrator has a root mean square of 8.06 and a median of 2.27, with only 28.7% of reactions within one reported standard deviation and 46.3% within two, so eQuilibrator understates its error. Group contribution, by contrast, has a median of 0.32 and 73.6% within one standard deviation, and dGPredictor performs even better with a root mean square of 1.15 and 87.9% within one standard deviation. The grading in the next section calibrates each source’s confidence against measurement rather than taking its uncertainty at face value.

### Direction assignment and evidence grading

Each source now generates its own prediction for the direction each reaction proceeds, and the three sources differ more in what they will commit to than in what they say (Figure 3A). eQuilibrator resolves 97.4% of the reactions it covers, group contribution 59.7%, and dGPredictor 40.1%. Across the database, 24,594 reactions receive a direction or a positive statement of reversibility from at least one source and 23,790 receive none.

**Figure 3.**
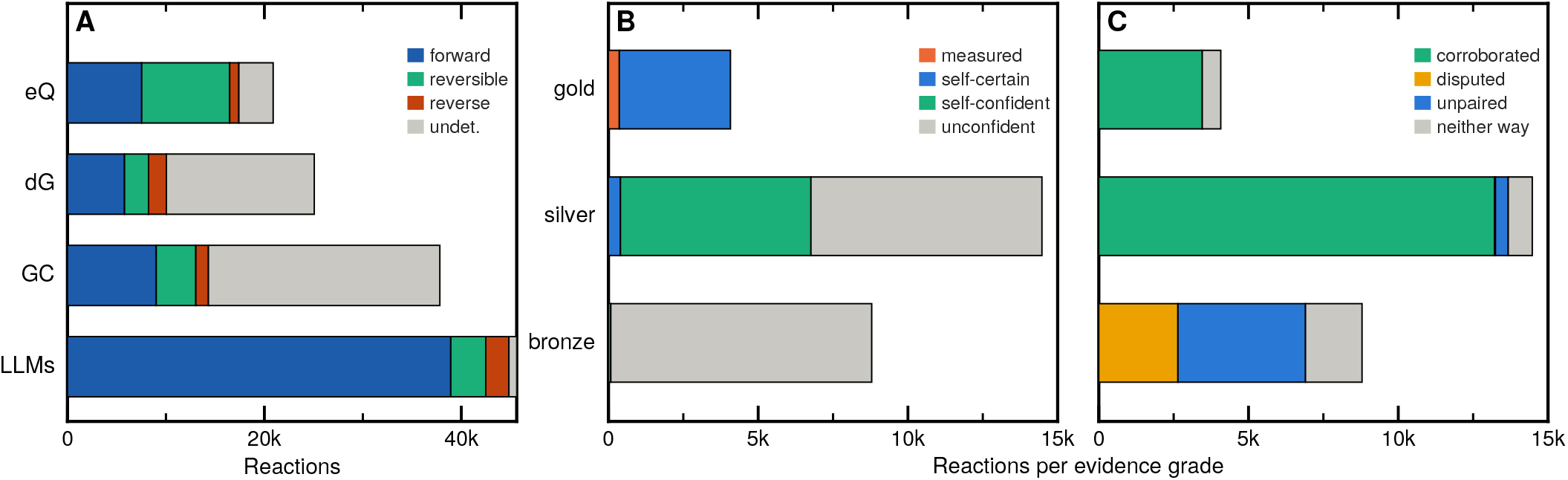
Direction, agreement, and evidence. (**A**) Direction assigned by each source independently: *eQ* eQuilibrator, *dG* dGPredictor, *GC* group contribution, and *LLMs* large language model ensemble that is not considered in evidence grading. Bar length is the number of reactions the source covers, facilitating comparison. (**B**) What each evidence grade rests on by the deciding source’s own calibrated confidence, and (**C**) by what the other sources made of it. A reaction is *self-certain* when the deciding source’s calibrated probability of being within 2.0 kcal mol^−1^ of the true value (*p*_ok_) is at *≥* 0.90, *self-confident* is 0.7 *≤ p*_ok_ *≤* 0.90, and unconfident is *p*_ok_ *≤* 0.7 (Supplementary Table 1).

#### eQuilibrator vs dGPredictor

dGPredictor is able to generate a predicted energy for more reactions (∼ 25, 100, against eQuilibrator’s ∼ 17, 900) and converts the least of that coverage into a usable reaction direction. For a typical reaction, the error bar from dGPredictor alone exceeds the decision threshold in the reversibility index, so 59.9% of its reactions are returned undetermined versus 2.6% from eQuilibrator. These sources agree 94.8% when they commit to a direction, although eQuilibrator provides more unique predictions (11,237) than dGPredictor provides (3,877) because dGPredictor often abstains rather than contradicting.

**Table 1.** shows how many reactions were graded as such, based on the evidence.

| Grade | Self-assessment | Cross-source | Parameters | Reactions | Conflict |
| --- | --- | --- | --- | --- | --- |
| gold | measured | — | stereo-exact anchor | 363 | 0 |
| gold | self-certain | — | $p_{\text{ok}} \geq 0.90$ | 3,710 | 14 |
| silver | self-certain | disputed | $p_{\text{ok}} \geq 0.90$; $R > 2$; $z > 2$ | 31 | 0 |
| silver | self-certain | unpaired | $p_{\text{ok}} \geq 0.90$; $n_{\text{src}} = 1$ | 370 | 0 |
| silver | self-confident | — | $0.70 \leq p_{\text{ok}} < 0.90$ | 6,361 | 22 |
| silver | unconfident | corroborated | $p_{\text{ok}} < 0.70$; $R \leq 2$; $z \leq 2$ | 7,713 | 0 |
| bronze | self-confident | disputed | $0.70 \leq p_{\text{ok}} < 0.90$; $R > 2$; $z > 2$ | 79 | 0 |
| bronze | unconfident | disputed | $p_{\text{ok}} < 0.70$; $R > 2$; $z > 2$ | 2,563 | 27 |
| bronze | unconfident | — | $p_{\text{ok}} < 0.70$ | 6,148 | 100 |
| **Total** |  |  |  | **27,338** | **163** |

#### Grading

Of 27,338 graded reactions, 4,073 are gold, 14,475 silver and 8,790 bronze (Figure 3B,C; see classification scheme in Supplementary Methods S3). The tiers separate on what supports them: gold is 91% self-certain and 85% corroborated while bronze is 99% unconfident, either uncorroborated (48%) or contradicted (29%) by other sources. Because a measured reaction is always graded gold, the grades must be recomputed with openTECR withheld before they can be tested against it. Re-graded on the predictors alone, the 363 distinct anchored reactions split into 172 gold, 186 silver and 5 bronze. The gold and silver reactions fall within 2 kcal mol^−1^ of measurement 93.0% and 89.2% of the time (Supplementary Methods S3), hence we recommend these tiers for inclusion in a metabolic model and reserve bronze for review rather than use in models.

#### Recommendation

We recommend the consensus among sources, but when sources disagree on reaction direction, we prioritize eQuilibrator followed by dGPredictor and then group contribution. Following this tiered approach, eQuilibrator supplies 16,985 recommended reaction directions, dGPredictor provides 7,935, and group contribution provides 2,055, the source for each recommendation being recorded.

### Ensemble LLMs predictions

The ensemble generated calls for ∼ 46, 000 reactions (Figure 3) and ∼ 23, 000 directions for reactions that no thermodynamic source committed on. However, 85% of its calls are simply the direction the equation is written in (Supplementary Methods S4), so we exclude these calls from evidence grading and report them as a separate, clearly-labelled layer. Neither the thermodynamics nor the ensemble approaches capture the context of metabolic pathways that can drive a reaction against its own standard-state energy as metabolite concentrations shift downstream (25).

### Atom mapping

Atom mappings are published for 26,242 reactions, 54% of the database: 19,809 clean, and 6,433 that required salvage (which are tagged). As the mappings can be used to span a metabolic reconstruction, reactions needing one repair are likely to carry others.

## Discussion

Unlike the 2020 release, this update adopts a multi-source approach rather than promoting a single thermodynamic value, publishing four sources that each have their own uncertainty and estimated direction, to capture source diversity. Grading against experiment enables us to validate the approach: A researcher can take the highest-graded source and know what the grade means. As this is a generalized reference database, we intentionally exclude organism-specific data; a choice affecting transport reactions and metabolic reconstruction that we discuss in Supplementary Methods S6. A current limitation is that roughly half the database lacks thermodynamic values, constrained by missing or incomplete chemical structures. For the remaining reactions, by explicitly reporting discrepancies among thermodynamic sources, we aim to better serve researchers who rely on standardized biochemistry for genome-scale reconstruction, flux analysis, pathway design, and predictive modeling.

## Data availability

The ModelSEED Biochemistry Database is available at https://github.com/ModelSEED/ModelSEEDDatabase under the Creative Commons Attribution License, as versioned flat files and JSON, and archived at Zenodo (https://doi.org/10.5281/zenodo.22782602). Data directly derived from KEGG and MetaCyc remain subject to the licences of those resources. It is also queryable through a Solr endpoint and browsable at https://modelseed.org/biochem. Released structures, thermodynamic values with per-source uncertainty, evidence grades, atom mappings and the low-confidence reaction flags are all in that repository. The scripts that regenerate every published value, including the eQuilibrator cache rebuild, are under Scripts/Thermodynamics/.

## Acknowledgements

R.T.G. acknowledges all the hard-working individuals who contributed their time voluntarily to transcribe and check the myriad of numbers and text to create the re-curated openTECR v1.0. The full list of contributors can be found at https://github.com/opentecr.

## Author contributions

Contributions follow the CRediT taxonomy. S.M.D.S.: conceptualization, methodology, software, formal analysis, data curation, visualization, writing – original draft, project administration. C.S.H.: conceptualization, funding acquisition, supervision. A.F., J.P.F., V.S. and C.T.: resources, data curation, investigation; J.P.F. also writing – original draft. J.N.E. and F.L.: resources, software (supporting). R.T.G.: the openTECR re-curation of the experimental measurements. E.N. and M.E.B.: eQuilibrator. V.U., M.A. and C.D.M.: dGPredictor. S.H. and Z.N.: atom mapping. A.P.A., R.W.C. and E.M.W.-C.: funding acquisition, supervision. All authors reviewed the manuscript.

## Notice of copyright

The submitted manuscript has been created by UChicago Argonne, LLC, Operator of Argonne National Laboratory (“Argonne”). Argonne, a U.S. Department of Energy Office of Science laboratory, is operated under Contract No. DE-AC02-06CH11357. The U.S. Government retains for itself, and others acting on its behalf, a paid-up nonexclusive, irrevocable worldwide license in said article to reproduce, prepare derivative works, distribute copies to the public, and perform publicly and display publicly, by or on behalf of the Government. The Department of Energy will provide public access to these results of federally sponsored research in accordance with the DOE Public Access Plan. http://energy.gov/downloads/doe-public-access-plan

## Supplementary Information

## S1 Structure curation

A compound’s structure is the foundation of the database: formula, charge, protonation, decomposition and thermodynamic estimation all depend on it. Where the source databases disagree, we have to choose, and the choice has to be recorded in a way that a researcher can follow.

### S1.1 Assignment

Structures currently arrive from four sources (MetaCyc, KEGG, ChEBI and Rhea), and one structure must be selected to represent a compound. We use the standardization of structure by InChI to allow us to find whether the set of structures assigned to a compound are identical or not. Frequently, the structures are not identical depending on the sources, for different reasons, and we attempt to resolve these manually. The reasons for our choices on a case-by-case basis are recorded in the repository, and many choices follow a convention, such as prioritizing a source (e.g. MetaCyc over KEGG) or stereochemistry, etc. ∼ 96% of the 35, 000+ compounds that are assigned a structure are done so without conflict, and many, but not all, of the remaining structures were manually resolved.

### S1.2 Conflict

There are compounds that still have conflicting structures, and we maintain two reports within the repository that are regenerated on every structure change and released with the database. `Structure_Conflicts.txt` carries ∼ 3, 000 structures for ∼ 1, 000 compounds that differ in any way; `Formula_Conflicts.txt` carries ∼ 140 structures across 56 compounds where the actual elemental composition of the structure differs. The manual selection of a structure to represent a compound does not override the presence of a structure in these two files, which allows the curation to be auditable.

### S1.3 Manual curation

A curator resolves a conflict by adding one row to a file of their own under `Biochemistry/Curation/overrides/structure_picks,` and opening a pull request within GitHub. One of the maintainers can run the set of scripts used to validate the selection and integrate the pick and its impact throughout the biochemistry database. The procedure is documented for contributors in `Structure_Curation_Overview.md` at the repository root. As the impact of any selection can be extensive, contributors are asked not to run the update pipeline themselves. This release carries 110 such picks, alongside 58 formula overrides for acyl-carrier proteins.

## S2 Thermodynamic conditions

### S2.1 Hydrogen

`equilibrator_api` ships one named condition set, its physiological default: pH 7.5, pMg 3.0, ionic strength 0.25 M, 298.15 K. We match it in three parameters, but report at pH=7 for continuity: The database and the modeling approaches adopted by the ModelSEED default to a pH of 7.

### S2.2 Magnesium

`equilibrator_cache` uses a default of pMg 10 (effectively magnesium-free) while requiring pH, ionic strength and temperature to be passed explicitly. We set pMg 3.0, about 1 mM free Mg^2+^, on the grounds that intracellular free magnesium is of order 0.5–2 mM in most cell types and that ATP is predominantly magnesium-bound *in vivo*. In the TECR corpus, nearly 3, 000 measurements (66%) record no magnesium concentration and the loader substitutes pMg 14. The 1,512 measurements that do record it have a median pMg of 2.4, about 4 mM. We recompute Δ_r_*G*^*′°*^ at pMg 14 and at the reported pMg 3.0 over a sample of 400 reactions containing at least one of the 510 magnesium-binding compounds in the cache; the median difference is 0.044 kcal mol^−1^. Across the whole database, ∼ 34, 400 of ∼ 56, 000 reactions (61%) contain a magnesium-binding participant and for most of them the shift is far inside the reported uncertainty. It is not negligible for a reaction whose direction call already sits near the reversibility threshold, but upon recomputation, we find that this only affects 50 reactions in the entire database, and if so, it only changes the direction between irreversible and reversible, never changing direction.

## S3 Evidence grading

Every reaction carrying at least one thermodynamic estimate is graded gold, silver or bronze according to the evidence available. We do not use the direction calls generated by the LLMs as described in the next section. We use a value of *τ* = 2.0 kcal mol^−1^ to claim that a heuristic is “correct” if it lands within *τ* of the reference value.

### S3.1 Calibration

The uncertainties (*σ*) reported by each source are not comparable: group contribution overstates its error by roughly 2.2× and eQuilibrator understates by 1.6×, so we don’t apply a threshold to *σ* directly. A probability is comparable in a way *σ* is not, so for each source we fit one as a function of *σ* by isotonic regression. The fit combines two kinds of observation: reactions judged against an experimental measurement, and reactions judged against a tight-uncertainty reference source, the former carrying three times the weight of the latter. Writing *e*_*s*_ for the error of source *s* on a reaction, the absolute difference between its estimate and the reference value, *e*_*s*_ = |Δ*G*_*s*_ − Δ*G*^*^|. What is fitted is the probability that this error stays within *τ*, given the uncertainty the source reported:

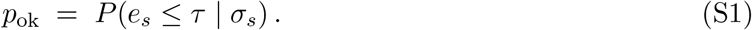

Running the same fit against the magnitude of the error, rather than an indicator of whether it cleared *τ*, gives a second calibrated quantity: the error a source is expected to make at a given *σ*,

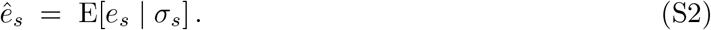

Both curves are fitted once per source. Applying either to a reaction requires only that source’s *σ*. Here we use *p*_ok_ to grade a reaction on its own evidence, and in the next section we used *ê*_*s*_ to grade a reaction on the agreement between sources.

### S3.2 Corroboration

The calibration allows us to ask whether a source is trustworthy on its own terms. Whether the sources *agree* is a different question. Here we use *ê*_*s*_ (S2) it is energy in kcal mol^−1^, as opposed to a dimensionless probability. The scale is floored at 0.3 kcal mol^−1^, since an *ê*_*s*_ of zero would carry infinite weight:

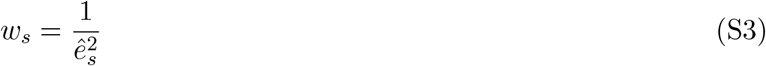

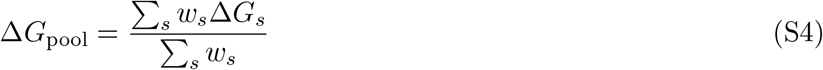

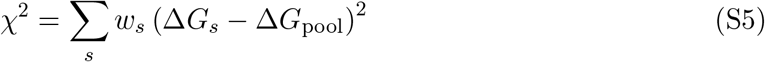

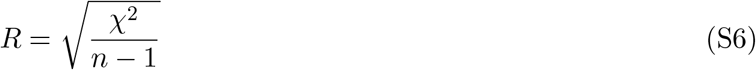

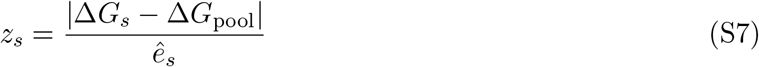

Taking these in turn: *w*_*s*_ (S3) is the precision of source *s*, so a source with a tighter calibrated error has more influence. Δ*G*_pool_ (S4) is the resulting weighted mean, an internal construct. *χ*^2^ (S5) measures how far the sources scatter about it, each deviation weighted in proportion to how confident that source claimed to be. *R* (S6) is that scatter per degree of freedom, where *n* is the number of sources contributing. *z*_*s*_ (S7) is the residual of a single source.

*R* compares the scatter among the sources with the uncertainties they themselves report. If reported uncertainties are accurate, the scatter among sources approximates *σ*, giving *R ≈* 1. Values well above 1 mean the sources lie further apart than their own error bars permit, so at least one is wrong. This quantity is partly what corroboration is decided on: *R* says whether the sources agree, and then *z*_*s*_ is used to determine which source is responsible when they do not.

### S3.3 Classification

Every reaction is graded as gold, silver, or bronze, and the grade is composed of two judgements: whether a source’s own calibrated confidence claims (*p*_*ok*_), and what the other sources make of it (*R, z*_*s*_). We enumerate here the rules we use to set the grade in roughly the descending order of authority:

1. **Measured**. A reaction carrying an experimental value is gold.
2. **Self-certain**. A reaction with *p*_ok_ *≥* 0.90 is gold, *≥* 0.70 is silver (self-confident), otherwise bronze (unconfident).
3. **Corroborated**. A reaction where *R ≤* 2.0 and *z*_*s*_ *≤* 2.0 will be promoted from bronze to silver.
4. **Disputed**. A reaction where *R* > 2.0 and *z*_*s*_ > 2.0 is demoted. The same pair of constants governs both a promotion and a demotion: a set is either concordant or it is not.
5. **Unpaired**. A reaction is demoted where it only has a single source of evidence.

Table 1: Reaction grades. A reaction takes the best grade any single source earns; ties are broken by the recommendation precedence (measurement, eQuilibrator, dGPredictor, group contribution), so the labels describe the strongest source rather than an arbitrary column order. *Conflict* counts reactions where eQuilibrator and dGPredictor both commit to an irreversible direction and those directions are opposed. Only 370 of the 4,686 reactions tagged `unpaired` appear in that row, as their grade is first defined by *p*_*ok*_. Counts are over non-obsolete reactions

## S4 AI assignment of reaction direction

The assignment of a reaction direction from thermodynamics means that nearly half the database doesn’t receive a direction ultimately due to the coverage of structures in the database, however well understood the reaction may be. In order to provide an alternative perspective of the reaction direction and to improve the coverage across more reactions in the database we implemented a complementary approach that predicts the direction of the reaction using an ensemble of large language models (LLMs). A prompt was formed that presented only the reaction identifier, its name, and its equation, with the equation written without a direction. None of the data generated from any of the heuristics or the experimental data from openTECR was used in the prompt.

### S4.1 Ensemble

Eleven candidate LLMs were scored across the role of proposer, auditor, and adjudicator (see next paragraph on how these are used). They were evaluated based on the rate at which an assigned reaction matched experiments. In the end, we selected: three separate LLMs from different commercial providers as predictors: Claude Opus 4.8, Gemini 3.5 Flash, and GPT-5.6; GPT-5.5 as the auditor and Claude Opus 5 as the adjudicator. The proposers were instructed to respond with a predicted direction (>, *<*, =, or ? where it judges the evidence insufficient), a confidence, and a written justification grounded in the stoichiometry or in known enzyme biology. The auditor compiled a vote table of the proposals and the adjudicator makes the final call.

### S4.2 Reactions and status

We filtered out reactions for which the prediction would not have been appropriate, this includes reactions flagged as obsolete, symmetrical transport reactions, and pseudo-reactions that acted as lumped reactions, aggregating many species and for which there is no direction to infer, leaving ∼ 46, 000 reactions for which direction is assigned. We store the output of the ensemble in our repository for review, including any objections raised by the auditor.

### S4.3 Summary of results

As the set of reactions covers almost all those for which we generated thermodynamics data, we’re able to draw a direct comparison. The ensemble commits to a direction far more readily than the thermodynamic rules do, abstaining on ∼ 2% of reactions, and it supplies a direction for 22,963 of the 23,790 reactions no thermodynamic source will commit on. With the far wider coverage, researchers may find the results useful, but we must note a caveat: 85.3% of the ensemble’s calls are the direction the equation is written in (>), against 5.2% reverse and 7.8% reversible. Reaction equations are typically written as if in the physiological direction, from left to right. This is a prior that exists and is not using thermodynamics. This is why the calls are released as their own layer and take no part in the evidence grading. But either a case where the ensemble calls a reverse direction, or where it disagrees with the thermodynamics evidence, may contribute information.

## S5 Where the growth since 2020 landed

As we did not add new KEGG biochemistry, and Rhea has no pathway ontology, we may only comment on the pathway-by-pathway addition of reactions from MetaCyc: of the 12,259 reactions added from MetaCyc, nearly ∼ 20% were assigned to a class of pathways: Supplementary Figure S1. The classes that grew are those of specialised metabolism and cell-envelope biosynthesis. Individual pathways that gained most include mycolate biosynthesis, arachidonate biosynthesis, colibactin biosynthesis and three branched-chain fatty-acid pathways. Conversely, central carbon and energy metabolism account for 41 of the 12,259 additions.

**Figure S1:**
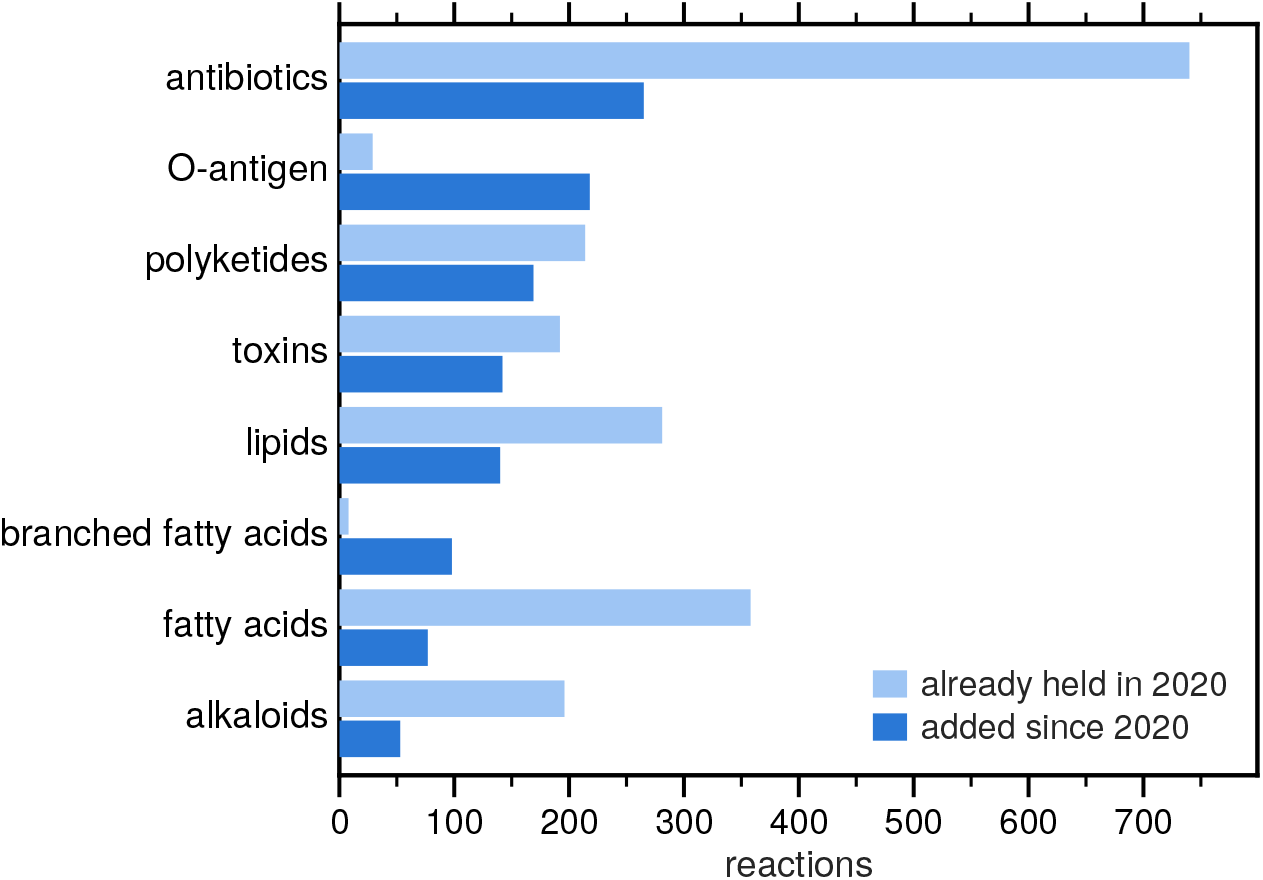
Where the growth since 2020 landed. Reactions from MetaCyc that were added since 2020 against those already held. Here we show the eight classes of pathways for which we added the most reactions. Growth is concentrated in specialised metabolism and cell-envelope biosynthesis.

## S6 A generalized reference database for ModelSEED

### A generalized reference database

ModelSEED biochemistry is the foundation of ModelSEED, but distinct from its automated pipeline for reconstruction of bacterial metabolism [1, 2], as well as its counterpart in flowering plant metabolism: PlantSEED [3, 4]. Due to the diversity of organisms generated by these pipelines, we strive to maintain an impartiality within this database that would otherwise affect how the models behave.

### Transport

Transport reactions are represented by casting each metabolite into separate compartments, but as many transport reactions may bridge different combinations of compartments, we use agnostic indices: 0 for one compartment and 1 for another. So one reaction can be instantiated against a bacterial membrane, a mitochondrion or a plastid envelope when ModelSEED or PlantSEED reconstructs a model. For this reason, the thermodynamic energy computed for a transport reaction is done so from stoichiometry alone and not from membrane potential or pH gradient.

### Reconstruction scope

The automated reconstruction pipelines in ModelSEED and PlantSEED involve the mapping of biochemistry to enzymatic annotation that is captured in the form of a template, which also includes the custom rules for the assignment of transport. Currently, the union of the ModelSEED v2 and PlantSEED v2 templates houses ∼ 9, 000 reactions but this is not a limitation on the scope of the database, and researchers may continue to use any reaction in our database as deemed useful.

